# High-throughput optical mapping of replicating DNA

**DOI:** 10.1101/239251

**Authors:** Francesco De Carli, Nikita Menezes, Wahiba Berrabah, Valérie Barbe, Auguste Genovesio, Olivier Hyrien

## Abstract

DNA replication is a crucial process for the universal ability of living organisms to reproduce. Existing methods to map replication genome-wide use large cell populations and therefore smooth out variability between chromosomal copies. Single-molecule methods may in principle reveal this variability. However, current methods remain refractory to automated molecule detection and measurements. Their low throughput has therefore precluded genome-wide analyses. Here, we have repurposed a commercial optical DNA mapping device, the Bionano Genomics Irys system, to map the replication signal of single DNA molecules onto genomic position at high throughput. Our methodology (HOMARD) combines fluorescent labelling of replication tracks and nicking endonuclease (NE) sites with DNA linearization in nanochannel arrays and dedicated image processing. We demonstrate the robustness of our approach by providing an ultra-high coverage (23,311 x) replication map of bacteriophage λ DNA in Xenopus egg extracts. HOMARD opens the way to genome-wide analysis of DNA replication at the single-molecule level.

## Introduction

DNA replication is a crucial biological process that ensures accurate transmission of genetic information to daughter cells^1^. Eukaryotic organisms replicate their genome from multiple start sites, termed replication origins, that are stochastically activated in S phase to establish bidirectional replication forks that progress along the template until converging forks merge^1–4^. Understanding the regulation of DNA replication is essential as perturbations of this process, referred to as replication stress, are today recognized as a major threat to genome stability and have been associated with cancer and other diseases^5^. However, mapping genome replication in metazoans has long remained challenging. For example, genome-wide mapping of human replication origins has been achieved by multiple approaches with only modest agreement^3,6^. One possible source of discrepancy is the incomplete purification of sequenced replication intermediates. A further limitation is that genome-wide replication mapping so far has been restricted to cell populations. As there is considerable cell-to-cell variability, ensemble profiles are at best a smoothed average of individual replication patterns. Single-molecule (SM) techniques must be used to fully evaluate cell-to-cell heterogeneity and uncover spatial correlations in replication patterns. SM techniques are also required to visualize rare pathological events such as fork stalling. Furthermore, SM replication signals are free from contaminating DNA molecules. Thus, the development of SM methods is crucial to progress in our understanding of DNA replication.

Unfortunately, none of the existing SM techniques has lend itself to full and robust automation. Consequently, their throughput has remained drastically low. Typically, cells are pulse-labelled with tagged nucleotide analogs, genomic DNA is purified, stretched on glass coverslips by DNA combing^7^ or other techniques^8^, and the tagged nucleotides are detected by several layers of antibodies^7,8^. The labelled DNA stretches reveal fork progression during the pulse and allow to infer origin positions and fork speeds, but contain no sequence information. This information can be obtained by hybridizing fluorescent DNA probes^9,10^, but only very few DNA molecules contain the probed locus. Months of work are required to collect and analyse a statistically significant sample of molecules for a single mammalian locus^11,12^.

Using replication-competent Xenopus egg extracts, we recently described a straightforward method for optical mapping of DNA replication (OMAR)^13^. Bacteriophage λ DNA was replicated in egg extracts in the presence of a fluorescent dUTP, purified, nicked with a site-specific nicking endonuclease (NE), nick-labelled with another fluorescent dUTP, stained with YOYO-1, and combed. Direct epifluorescence revealed, in three distinct colours, the DNA molecules, their replication tracts and their NE sites, allowing alignment to a reference genome^13^. The irregular surface deposition of combed DNA molecules, nevertheless, remained refractory to robust automated analysis. However, elongation of DNA in parallel nanochannel arrays has recently emerged as a potential alternative to DNA combing^14^ and a commercial automated platform, the Bionano Genomics Irys system, demonstrated ability to assemble large numbers of nick-labelled DNA molecules into genomic maps^15,16^.

Here, we have exploited the Irys system for high-throughput optical mapping of replicating DNA (HOMARD) using the same fluorescent labelling strategy as in OMAR. We developed all necessary computational tools to robustly extract replication signal with high throughput. We report the high-quality imaging, detection and mapping of 43,493 barcoded, replicating DNA molecules in Xenopus egg extracts. The results demonstrate the feasibility of high-throughput mapping of DNA replication at the SM level and provide an ultra-high (23,311 x) replication coverage of *in vitro* reconstituted λ-DNA minichromosomes, revealing replication landscape with groundbreaking precision.

## Methods

### Preparation of labelled DNA replication intermediates for nanochannel analysis

Replication of λ DNA in the presence of 20 μM AlexaFluor647-aha-dUTP (AF647-dUTP) in Xenopus egg extracts and purification of the DNA in agarose plugs was as described previously^13^, except that pelleted nuclei from 5 × 50 μl replication reactions were pooled together in a single agarose plug and that proteinase K digestion was performed within a 5-fold larger volume of the same digestion buffer. Nicking at Nt.BspQI sites, nick-translation with AF546-dUTP, YOYO-1 DNA staining and sample run was performed according to Bionano Genomics specifications for the Irys system.

### Data collection

DNA molecules were linearized and imaged using the Bionano Genomics Irys system. Each Irys nanofluidic chip contains two flow cells, ~13,000 nanochannels each. DNA molecules are driven by electrophoresis into the nanochannels where they are rectilinearly stretched by physical confinement. Molecules are held still by stopping electrophoresis, automatically imaged (1,140 images per scan) and electrophoresis is resumed to image new DNA molecules. The cycle is repeated 30 times yielding a total 34,200 images per run per colour. DNA molecules (outlined by YOYO-1 staining) and locations of fluorescent nick-labels along each molecule are detected using the AutoDetect software (v 2.1.4.9159) provided by Bionano Genomics.

### Illumination and chromatic shift corrections

To correct raw images for inhomogeneous illumination, single pixel median of red, green and blue images were computed for each of the 30 scans composing a run and subtracted from each of the 1,140 images of the same scan. To correct for chromatic shift, we selected the 15% brightest, illumination-corrected images of each scan (171 out of 1,140 scan images) which are enriched in foreground (i.e. molecules) and thus more informative. From this image subset (5,130 out of 34,200 run images) we extracted sets of points corresponding to local maxima for each of the three channels and paired the closest points for both the red/blue and the green/blue pairs. Then, we determined for each image and each colour pair the affine transformation matrix that minimized the distances between the paired points. Matrix parameters clustered around very similar median values for each scan of a run. We therefore computed the median parameter values of the total run to define a unique transformation matrix that was then applied to the 34,200 images of the run.

### Optical mapping

To map λ DNA concatamers, we built a synthetic reference genome composed of 30 λ DNA genomes in head-to-tail concatamer (48,502 bp each, 1,455,060 bp in total). The fasta genome was *in silico* digested to Bionano cmap using Knickers software (v 1.5.5). Optical mapping was performed using the IrysView Genomic Analysis Viewer (v 2.5.1.29842) and RefAligner (r5122) from Bionano Genomics with the following parameters:

~~~
-nosplit 2 -BestRef 1 -biaswt 0 -Mfast 0 -FP 1.5 -FN 0.15 -sf 0.2 -sd 0.0 -A 5 -
outlier 1e-3 -outlierMax 40 -endoutlier 1e-4 -S -1000 -sr 0.03 -se 0.2 -MaxSF 0.25 -
MaxSE 0.5 -resbias 4 64 -maxmem 27 -M 3 3 -minlen 90 -T 1e-7 -maxthreads 4 -hashgen
5 3 2.4 1.5 0.05 5.0 1 1 3 -hash -hashdelta 10 -hashoffset 1 -hashmaxmem 64 -
insertThreads 6 -maptype 0 -PVres 2 -PVendoutlier -AlignRes 2.0 -rres 0.9 -
resEstimate -ScanScaling 2 -RepeatMask 5 0.01 -RepeatRec 0.7 0.6 1.4 -maxEnd 50 -
usecolor 1 -stdout -stderr.
~~~

Only DNA molecules ≥90 kb with label signal-to-noise ratio (SNR) ≥2.75 (static) were used for mapping.

### Intensity profile extraction

Once the three colour channels were corrected for colour shifts, the intensity profiles along the length of each molecule were automatically extracted using both X and Y molecule coordinates identified by Autodetect (Xstart, Ystart - Xend, Yend) and a 3 px width-window, in order to correct for the observed slight tilting of molecules with respect to image borders and to fully enclose each molecule’s contour without overlapping adjacent molecules. Results were exported in a plain table (MoleculeIntensityProfile.csv) and appended to the original Molecules.mol Bionano file for further use.

### Alignment of single-molecule intensity profiles to the reference genome

To position SM intensity profiles to their genomic location we linked information from several Bionano Genomics’ output files to intensity profiles contained in MoleculeIntensityProfile.csv produced at the previous step. We recovered the ID of the first correctly mapped molecule’s nick label (*FirstLabel_ID*) and its genomic position (*GenomPosStart*) from the MoleculeQualityReport.xmap file generated after optical mapping and its distance from the molecule start from the Labels.lab file *(FirstLabel_Position)* generated by Autodetect and corrected for chromatic shift. We then recovered genomic coordinates of all molecule intensity profiles by aligning the corrected *FirstLabel_Position* onto the *GenomPosStart*.

### Statistical testing

To test if the local mean replication signal and AT content of replicating λ-DNA monomers were linearly related to each other, the AT content profile of the λ genome was computed with a 533 bp sliding window (estimated number of bp per pixel) and the following statistical test was devised. As per the central limit theorem, the sample size being very large, the mean of the replicative signal follows a Gaussian distribution at each position. A Gaussian confidence interval at 1% risk was then obtained from the estimated mean and standard deviation of the sample mean at each position. This confidence interval was subsequently adjusted for multiple testing with the Bonferroni correction.

### Data availability

The data that support the findings of this study are available from the corresponding authors upon reasonable request.

## Results

### Linearizing and imaging DNA replication intermediates in nanochannel arrays

The experimental strategy for high-throughput visualization of nick-labelled DNA replication intermediates is shown on Figure 1A. Bacteriophage λ DNA (48.5 kb) was replicated in Xenopus egg extracts in the presence of AF647-dUTP (red), purified, nick-labelled at Nt.BstQI sites with AF546-dUTP (green), stained with YOYO-1 (blue) and run on a Bionano Genomics IrysChip. Automated imaging yielded 34,200 fields-of-view (FOVs) in 3 colours each per sample. A single FOV spans ~140 nanochannels of equivalent length to ~270 kb of B-form DNA stretched to ~85% of crystal length. Molecules of up to ~3 Mb may be imaged through 12 consecutively stitched FOVs.

**Figure 1.**
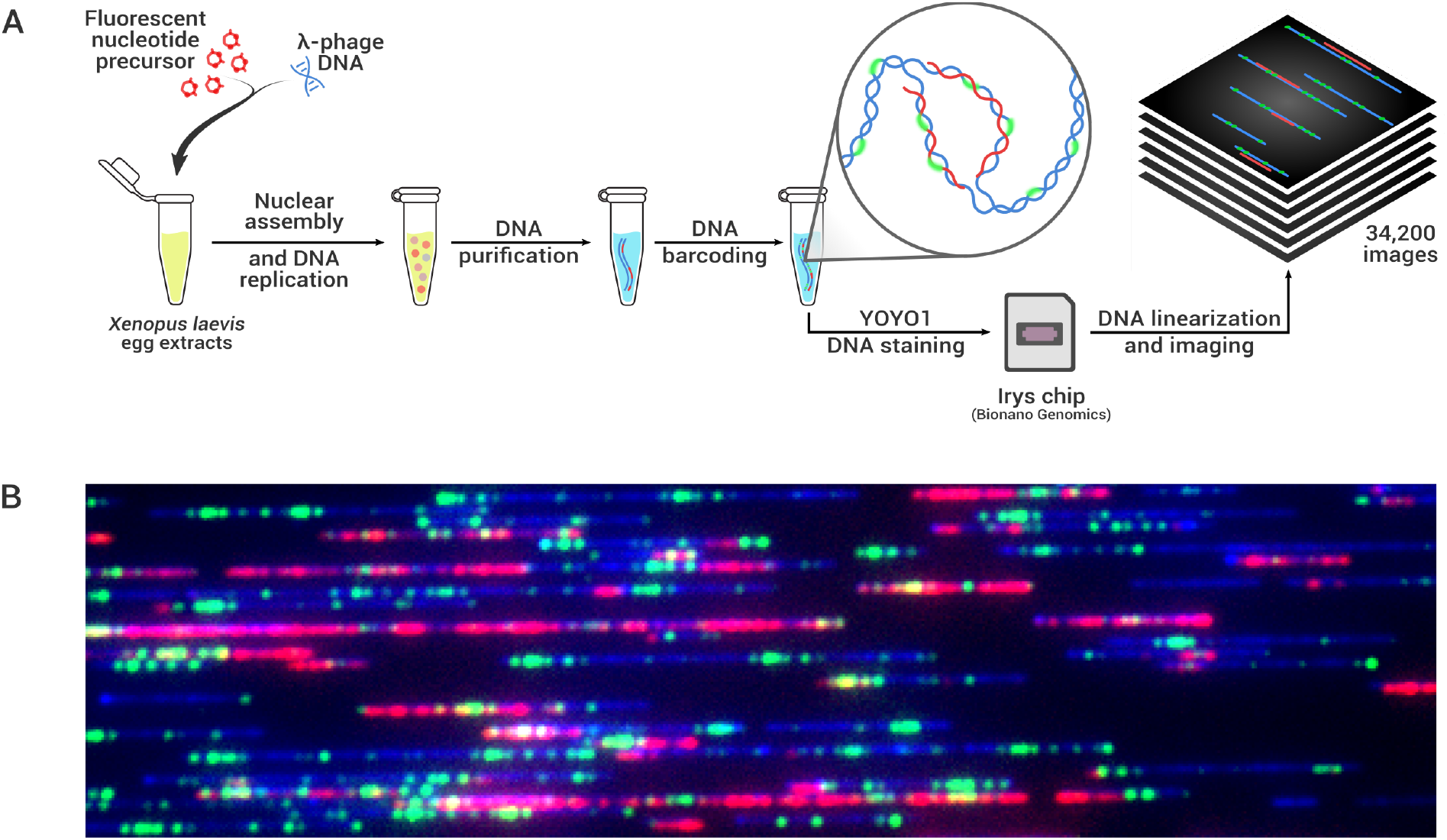
Experimental Approach. (**A**) Bacteriophage λ DNA (48.5 kb) was replicated for 3 h in *Xenopus laevis* egg extracts supplemented with AF647-dUTP (red). Purified DNA was nick-labelled at Nt.BspQI sites with AF546-dUTP (green), stained with YOYO-1 (blue), stretched in nanochannels and imaged using the Irys system (Bionano Genomics Inc.) yielding 34,200 FOVs containing a total of 63,890 Mb of DNA molecules (28,860 Mb in molecules >90 kb). (B) An exemplary merged image of stretched, 3-colour fluorescent DNA molecules spanning an entire FOV (270 kb) in the direction of stretching.

A typical FOV section (Figure 1B) shows a mixture of unreplicated and replicating DNA molecules. A broad distribution of molecular sizes was observed, because linear λ-DNA monomers are actively end-joined in concatamers in egg extracts^13^. Replicating DNA molecules showed alternating replicated and unreplicated DNA stretches 5–50 kb in size, as previously observed by DNA combing^13^. Compared to DNA combing^13^ or to earlier nanochannel experiments^14^, the advantages of the Irys system were the robust and highly uniform linearization of DNA, the lower background and the smoother signals, allowing high-throughput data collection and analysis.

### A robust pipeline for high-throughput replication tract imaging

The standard Irys visualization pipeline is shown on Figure 2A. Raw images are analysed with Autodetect to get DNA molecules’ coordinates (start; end) and barcode label positions. Schematized molecules are aligned to the reference genome using IrysView and RefAligner. Standard genome mapping applications of the Irys system only require two-colour images (DNA, blue + barcode, green), although the third colour (red) has been occasionally used for dual red/green nick-labelling experiments^17^.

**Figure 2.**
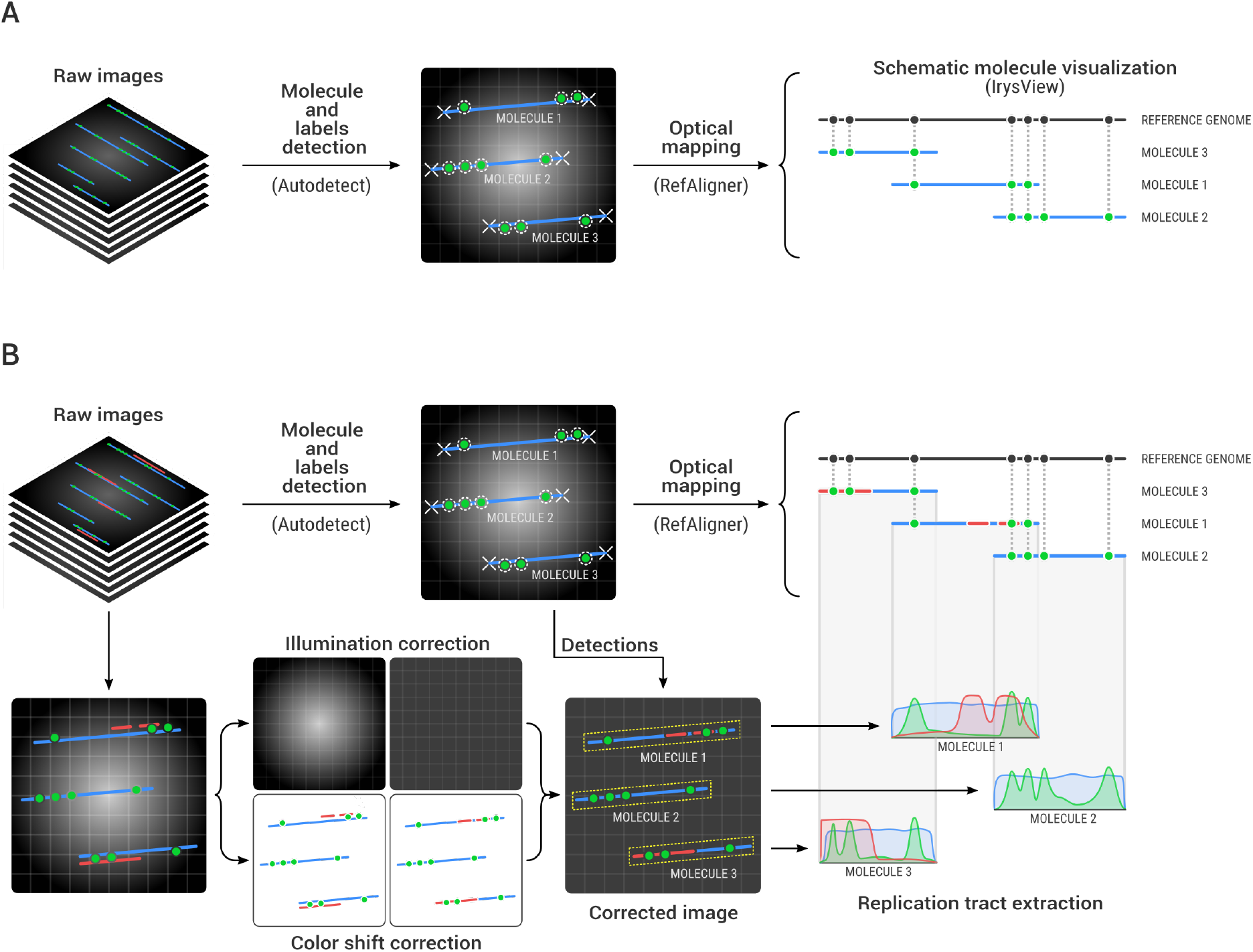
Bioinformatics for High-throughput Optical Mapping of Replicating DNA (*HOMARD*). **(A)** The current workflow for Irys data analysis and visualization consists of 3 steps. First, the DNA molecules and their labels (barcode) are automatically detected using Autodetect; second, individual barcoded DNA molecules are aligned to the reference genome using RefAligner; third, schematic results are visualised using IrysView. **(B)** A custom pipeline was developed to output SM intensity profiles (bottom pipeline). Raw 3-colour channels images (34,200 per run) are corrected for chromatic shifts between the three channels (blue = DNA, red = replication, green = barcode) and for illumination inhomogeneities. Corrected images display red replication tracts and green NE labels re-aligned onto their original DNA molecule (blue) over a homogeneous low background. Then, the Autodetect and RefAligner coordinates of each molecule are used to automatically extract the three colour intensity profiles and align them to the reference genome.

Here we took advantage of the third channel’s availability to image replication tracks and we developed a dedicated pipeline for HOMARD grounded on the original Bionano pipeline (Figure 2B). After DNA molecules and barcodes were detected and mapped using Autodetect and RefAligner, custom Python scripts were developed to correct images for inhomogeneous illumination and for chromatic shifts as well as to extract 3-colour intensity profiles for individual DNA molecules at high accuracy and throughput.

The raw images presented systematic inhomogeneous illumination with stronger intensity values at their centre. This effect was particularly clear when computing the median at each location (x,y) of the blue, red and green image channel of each scan (Figure S1). Those median images per scan showed channel-specific intensity ranges and shapes. For each scan, the blue channel had lower pixel intensities and a flatter shape than the red and green ones. As median images slightly varied between scans, we corrected the raw images by subtracting the median image of the corresponding scan, which successfully flattened images of the entire run.

Once the illumination biases were corrected, the composite images produced by the Irys system revealed a significant chromatic shift. The blue (DNA), green (barcode) and red (replication) signals of each molecule were not exactly aligned to each other (Figure 3). The misalignments were low at image centre but gradually increased towards the image borders and were more pronounced between red and blue than between green and blue. This observation was consistent with transverse chromatic shift. As molecules were automatically detected on the blue (YOYO-1) channel, it was necessary to rectify the coordinates of the red (AF647) and green (barcode) pixels in order to assign them to the correct DNA molecule. To align the channels, we used an affine registration approach were channels were assumed to be related to one another by an affine transform. Coefficients of a transformation matrix between channels were identified as minimizing the sum of squared error of a pair of feature point sets (see Methods). Such an affine transform comprises all global distortions that preserve lines, such as magnification, rotation and shift. The misalignment was satisfactorily corrected for most image pairs as shown in Figure 3.

**Figure 3.**
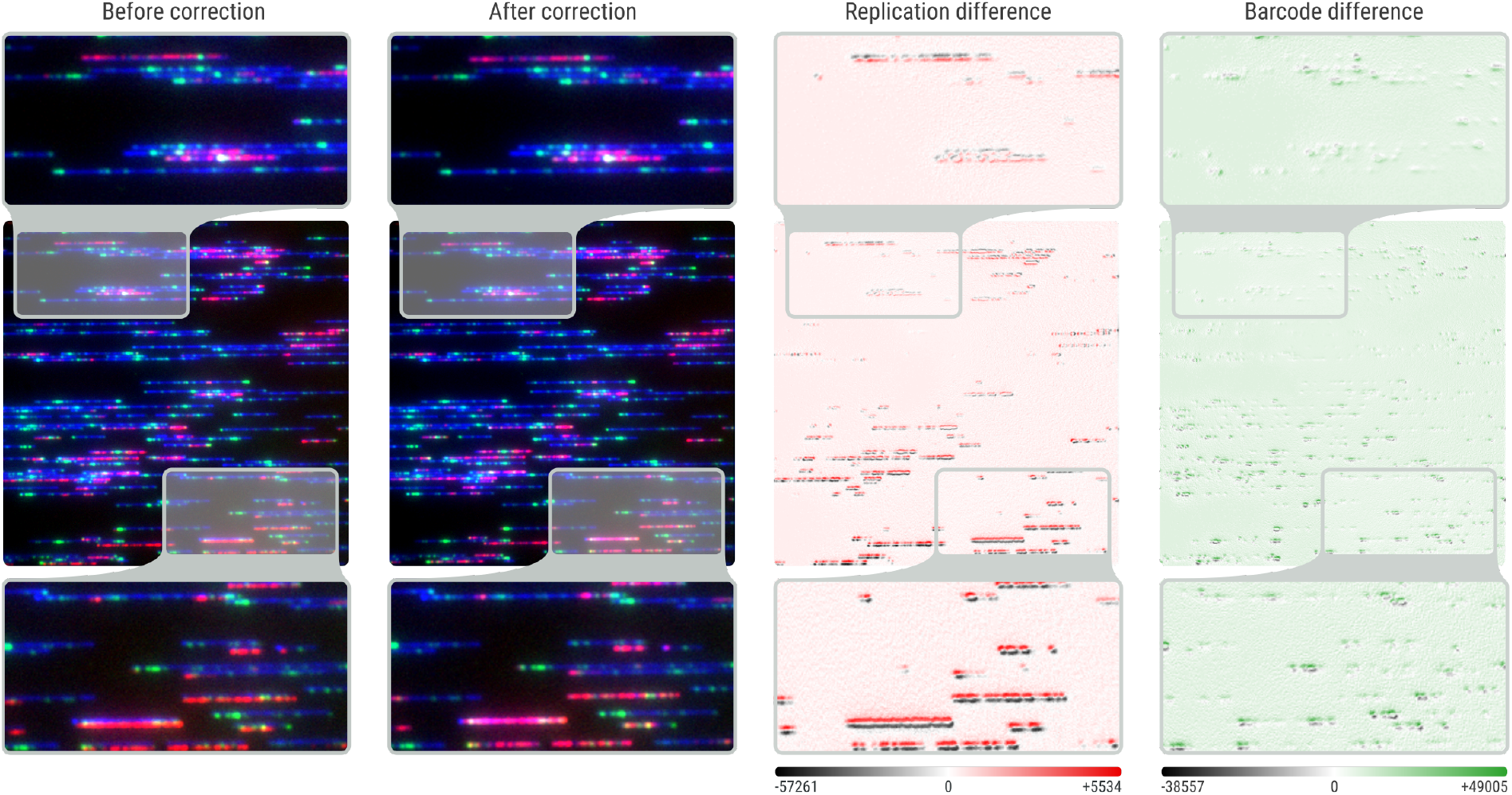
Chromatic shift correction. Raw 3-colour images present chromatic shifts between the red, green and blue channels. As a result, replication and barcode signals are shifted with respect to the DNA signal (Before correction). For all 34,200 images of a run, red and green images are automatically corrected to match the unaltered blue image (After correction). Chromatic shifts occur on both X and Y axes, and are stronger at image borders (zoomed-in molecules inside grey rectangles). Subtracting the uncorrected from the corrected red (Replication difference) and green (Barcode difference) images highlights the spatial chromatic shift gradient.

### High-throughput mapping and profiling of replicating DNA molecules

Once all images were corrected for colour shift and illumination biases, we used (1) the molecule coordinates provided by Autodetect to extract SM intensity profiles from them (taking into account the tilting of DNA molecules) and (2) the optical mapping results from RefAligner to align them onto their genomic coordinate (see Methods).

Autodetect reported 884,832 DNA molecules ≥ 20 kb totalling 63,245 Mb. Only DNA molecules ≥90 kb with label SNR ≥2.75 (static) were used for mapping (210,147 molecules totalling 28,860 Mb). Because λ-DNA monomers are end-joined in complex concatamers in egg extracts^13^, we built a synthetic reference genome composed of 30 λ-DNA molecules in a head-to-tail tandem array (1,455 Kb). Using RefAligner, 33.1% (N = 69,203) of the selected 209,051 molecules were mapped to the synthetic genome. The remaining molecules corresponded to more complex concatamers containing head-to-head and tail-to-tail junctions between λ-DNA monomers and/or incomplete monomers. For this report, we focused on molecules entirely contained in a single FOV (N = 120,793), of which 36.0% were mapped (N = 43,493).

Similar stretching of replicated and unreplicated segments is mandatory for unbiased mapping and analysis of replication intermediates. To check if any bias could have occurred at this level, we partitioned the 120,793 single-FOV molecules into unreplicated (N = 90,831) and replicating (N = 29,962) molecules based on their computed mean red signal intensity. The two datasets showed similar molecule size distributions and barcode label densities. Replicating intermediates were mapped to their *in silico* reference genome with only slightly lower efficiency (29.4%) than unreplicated molecules (38.2%). Therefore, 1) AF647-labelled tracts did not prevent nick-labelling with AF546-dUTP; 2) replication forks did not prevent DNA entry into nanochannels; 3) the close juxtaposition of sister duplexes in nanochannels did not result in different stretching of replicated and unreplicated DNA segments.

Figure 4 shows intensity profile extraction and genome positioning of an exemplary, 120 kb-long replicative DNA molecule. The raw image presented a downward shift of the red and green signals relative to the blue signal. Chromatic shift correction realigned the signals which increased the signal-to-noise ratio of the red and green intensity profiles. Illumination correction lowered the baseline of the three profiles, due to subtraction of the median background image, which further increased the signal-to-noise ratios. Note that without chromatic correction, the rightmost replication bubble on Figure 4 would have been missed. A complementary example of false signal collection from an adjacent DNA molecule in the absence of chromatic shift correction is shown on Figure S2. The molecule’s profiles were aligned to the reference genome by RefAligner based on the corrected green profile. Note that the green profile confirms the head-to-tail tandem orientation of two λ-DNA monomers. The two monomers were identically stretched despite the presence of replication bubbles on the first monomer but not the second one (Figure 4).

**Figure 4.**
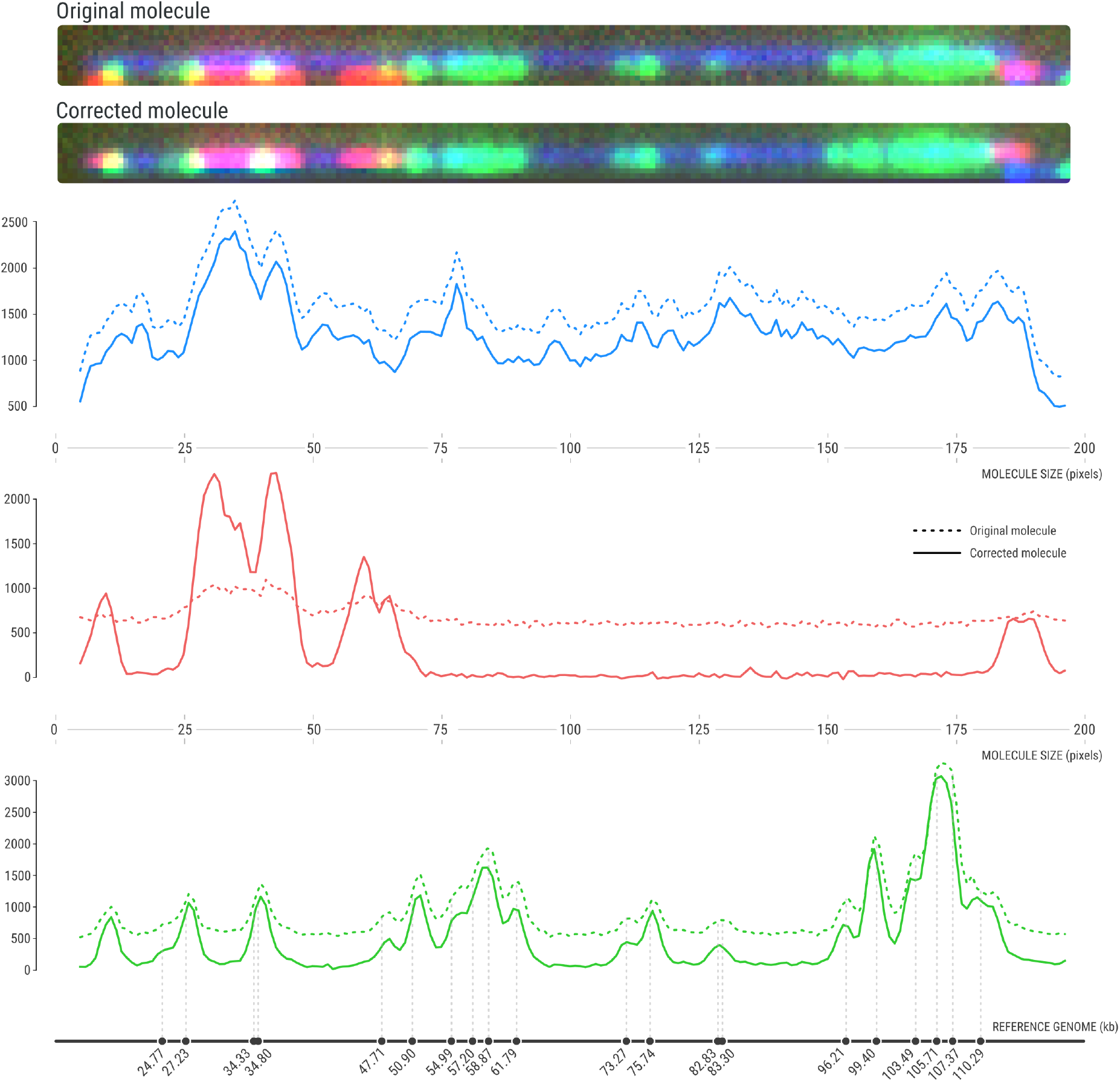
Single-molecule intensity profiles from a replicative DNA molecule. Image crops and fluorescent intensity profiles (blue, DNA; red, replication; green, barcodes) of a representative replicative DNA molecule before (top image, dotted line profiles) and after (bottom image, solid line profiles) chromatic shift and illumination correction. The length of the molecule is indicated in pixels under the blue and red profile. The reference genome coordinate is indicated in Kb under the green profile, with vertical dashed lines connecting the green profile peaks with their *in silico* map position.

We then created a software to simultaneously visualize the colour-coded intensity profiles of all mapped molecules. Figure 5 shows the full collection of 7,019 replicating λ-DNA concatamers mapped to (and included in) their synthetic reference genome and ordered by molecule start position. The correct genome alignment of the concatamers is visible from the repetitive pattern of vertical lines in the green channel (barcode). Half of the DNA molecules mapped to the first 7 λ-DNA monomers of the synthetic genome and all the molecules mapped to the first 14 monomers. Enlarged views of the aligned molecules illustrate, as expected, their continuous DNA signal (blue), discontinuous and irregular replication signal (red), and repetitive barcode signal (green) aligned with Nt.BstQI sites on the synthetic genome map.

**Figure 5.**
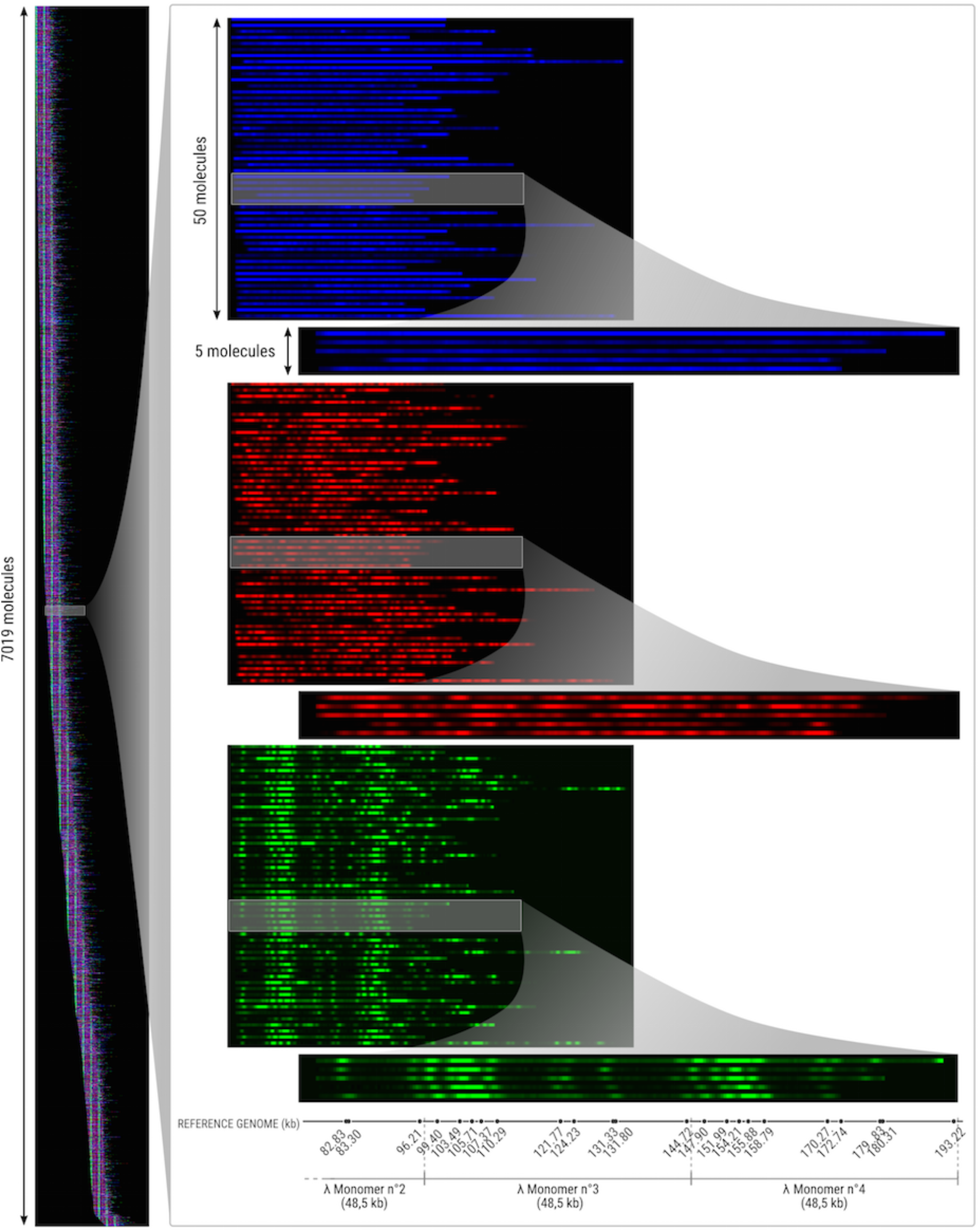
High-throughput, single-molecule replication analysis. **Left**, for each of the 7,019 replicating λ-DNA concatamers mapped to the synthetic, head-to-tail concatameric λ genome, the intensity signals collected from the three channels were used to create a horizontal, synthetic tricolour profile of 3 pixels height. The 7,019 tricolour profiles were mapped to their genomic location and vertically ranked by molecule start position. Blue, DNA; red, replication; green, barcodes. A repetitive pattern of green (barcode) vertical lines is visible, as expected for the successful optical mapping of λ-DNA concatamers. All molecules were mapped to the first 14 monomers of the 30 monomer-long synthetic genome. **Right**, consecutive zoomed-in views of selected regions indicated by shaded rectangles, with the three colours separately shown. The corresponding section of the synthetic genome map is aligned below the enlarged green molecule images. Nt.BstQI sites are indicated by black dots. Numbers below each dot indicate their map position on the synthetic λ-DNA concatamer genome.

### Sequence-dependence of λ-DNA replication signal

Several studies reported long ago that replication initiation is independent of DNA sequence in Xenopus eggs, egg extracts, and early embryos^18^. We recently confirmed this conclusion by OMAR analysis of λ-DNA in egg extracts^13^. Here, the mapping of 7,019 replicating λ-DNA concatamers, totalling 23,311 λ-DNA monomers, allowed us to reinvestigate this question with unprecedented precision. Since AF647-dUTP incorporation only occurs opposite to adenines on the DNA template, the signal from replicated tracts, if they are evenly distributed along the template, was expected to be strictly proportional to the local AT-content. Accordingly, we found that the profile of the mean red signal intensity of the 23,311 λ-DNA monomers closely followed the AT content profile (Figure 6: Pearson correlation *R* = 89%). Only small (<10%), albeit statistically significant (*P*<10^−8^) deviations were observed between the two profiles. While this result confirms the lack of strong sequence preference previously observed with less precise methods^18^, it also raises the possibility that replication tracks are not as uniformly distributed as previously thought. Further analysis will reveal if these small deviations reflect a subtle sequence dependence of λ-DNA replication initiation or elongation in Xenopus egg extracts.

**Figure 6.**
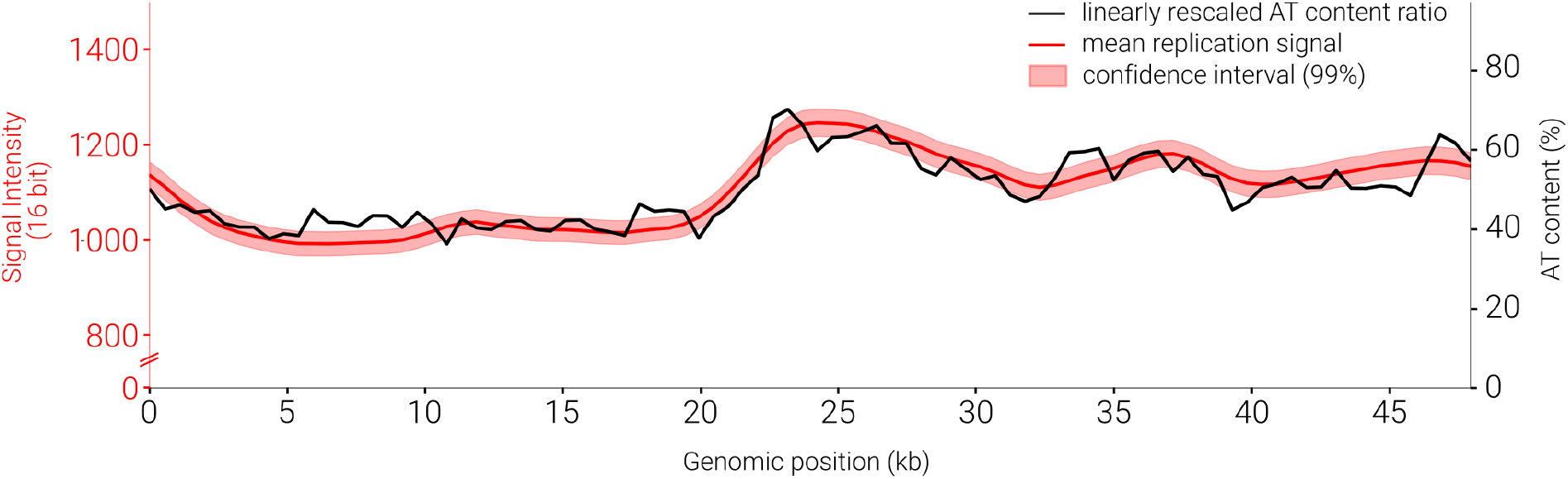
The mean replication signal of replicating λ-DNA molecules closely follows the local AT-content. Black line, AT-content of the λ-DNA sequence computed by non-overlapping 537 bp windows. Thin red line, mean replication signal (on a 16-bit scale) of 23,311 λ-DNA monomers from 7,019 replicating concatamers. Thick red line, 99% confidence interval after Bonferroni correction.

## Discussion

Despite the importance of SM information for understanding DNA replication, current SM replication mapping methods have remained refractory to robust automation and are therefore low throughput^11,12^. We recently described a new method for optical mapping of DNA replication (OMAR) that combines triple-colour fluorescent labelling of DNA, replication tracks and nicking endonuclease sites with DNA combing^13^. This method considerably simplified the labelling and mapping of DNA replication intermediates with respect to standard, antibody-based DNA combing, but remained difficult to automate. In this work, we leveraged the potential of OMAR replication labelling strategy by combining it with the Bionano Genomics Irys system’s power to automatically image and robustly map nick-labelled DNA molecules at high throughput. Compared to standard SM replication mapping techniques, HOMARD drastically accelerated data collection (from several months to a few days) and enormously increased dataset size (from a few hundreds to > 200,000 molecules in a single experiment). This was made possible thanks to the superior DNA linearization and imaging capacities as well as the dedicated tools, either provided by the Irys system for molecule detection and mapping, or additionally developed here for uneven illumination and chromatic shift correction and for extraction and alignment of replication signals.

Importantly, we demonstrate that DNA replication intermediates can be driven and extended into Irys nanochannel arrays with comparable efficiency to linear, nonreplicating DNA molecules. Furthermore, we observe a similar stretching of unreplicated and replicated DNA along replication intermediates, which was mandatory to map them based on distance measurements between consecutive nick-labels. Thus, chasing replication forks following replicative track labelling was not required to allow proper entry and stretching of DNA into nanochannels.

The standard Bionano Genomics pipeline can be used to detect DNA replication intermediates and their nick labels but is not suited to detect labelled replication tracks. Although the Autodetect software can detect both dotty (green nick labels) and elongated (blue DNA molecules) objects, our attempts to detect replication tracts using either option were unsatisfactory. Small tracts were missed when using the molecule’s detection option. The multiple red dots detected using the nick-label option did not accurately represent the true extension of replicative signals. We therefore developed scripts to correct for illumination defects and chromatic shifts, and to automatically output tricolour intensity profiles of multiple molecules aligned to their genomic map. In doing so, we found that image correction prior to intensity profile extraction increased the SNR and avoided signal omission or erroneous collection from adjacent molecules. Importantly, our pipeline is not limited to DNA replication but can be used for any type of epigenetic information that can be converted into a fluorescent signal, e.g. cytosine methylation^19^. Another advantage of our approach is to compact the useful replicative or epigenetic signal into one-dimensional intensity profiles, facilitating further analyses compared to direct feature detection on two-dimensional images.

Purified DNA added to Xenopus egg extracts is chromatinized and replicated in a manner that extraordinarily faithfully mimics DNA replication in the early embryo^20^. Taking advantage of this system, we produced an ultra-high coverage (23,311 x) singlemolecule replication map of eukaryotic minichromosomes made of tandem λ-DNA concatamers. The mean replicative signal from these minichromosomes was correlated with unprecedented precision to their AT-content, suggesting potential for optical sequencing of single DNA molecules. The lack of large deviations between the AT content and replication profiles is consistent with the lack of obvious sequence preference for replication initiation observed in previous studies in this system^13,18^. Further analyses will reveal whether the small deviations yet observed reflect a subtle sequence dependence of replication initiation or elongation and will explore potential functional correlations between neighbouring origins and forks.

HOMARD requires fluorescent metabolic labelling of replication forks, which is convenient to perform in Xenopus egg extracts by direct addition of fluorescent dNTPs. Permeabilisation procedures allowing fluorescent dNTP entry into mammalian cells have been developed^21^, which should extend the range of experimental systems accessible to HOMARD. Furthermore, a human cell-free system supporting fluorescent nucleotide labelling has been extensively used for DNA replication analysis^22–24^. In contrast, the use of 5-ethynyl-2’-deoxyuridine, which allows fluorescent labelling of newly synthesised DNA by click-chemistry, was reported to generate an unacceptable level of DNA breakage for isolation of long DNA molecules^14^. When working with unsynchronized cells, a dual-colour labelling scheme is necessary to orient replication forks and distinguish elongation from initiation and termination tracts. Unfortunately, the Irys system is currently limited to three fluorescent channels. Since two channels are required for DNA and nick label detection, this leaves only one channel for replicative labelling. One possibility is to vary the label concentration during the pulse and distinguish strong from weak tracts, as historically developed to demonstrate bidirectional replication by DNA fiber autoradiography in E. coli^25^ and in mammalian cells^26^. Alternatively, one could distinguish replicated from unreplicated DNA segments by their different YOYO-1 fluorescent intensity^13,27^ and use this information to orient the fluorescent dNMP tracks located at the ends of replicated segments. Both possibilities will necessitate dedicated automated analysis tools that we plan to develop.

In summary, this work demonstrates the potential of the Irys system for DNA replication and other functional genomic studies beyond its standard use for genome assembly and structural variation analysis.

## Acknowledgments

We thank Erwan Denis, Ghislaine Magdelenat and Caroline Belser (Génoscope) for their contribution to the production and analysis of optical mapping data, Felipe Delestro (IBENS) for help with Figure preparation, Gaël Millot (Institut Pasteur), Benoît le Tallec (IBENS) and other members of the O.H. lab for helpful discussions. This work was supported by the Ligue Nationale Contre le Cancer (Comité de Paris), the Association pour la Recherche sur le Cancer, the Agence Nationale de la Recherche (ANR-15-CE12-0011-01), the Fondation pour la Recherche Médicale (FRM DEI201512344404), the Cancéropôle Ile-de-France and the INCa (PL-BIO), the programme “Investissements d’Avenir” launched by the French Government and implemented by the ANR (ANR-10-LABX-54 MEMOLIFE and ANR-10-IDEX-0001-02 PSL*Research University) and the France Génomique national infrastructure, funded as part of the « Investissements d’Avenir » program (ANR-10-INBS-09).

## Author Contributions

**Experiments**: FdC, WB. **Software**: NM. **Data Analysis**: NM, FdC, AG, OH. **Project Supervision**: VB, AG, OH. **Writing-Draft**: FdC, NM, OH. **Writing-Editing**: FdC, NM, AG, OH. **Project Conception, Funding Acquisition, Administration**: OH.

## Supplementary Figures

**Figure S1.**
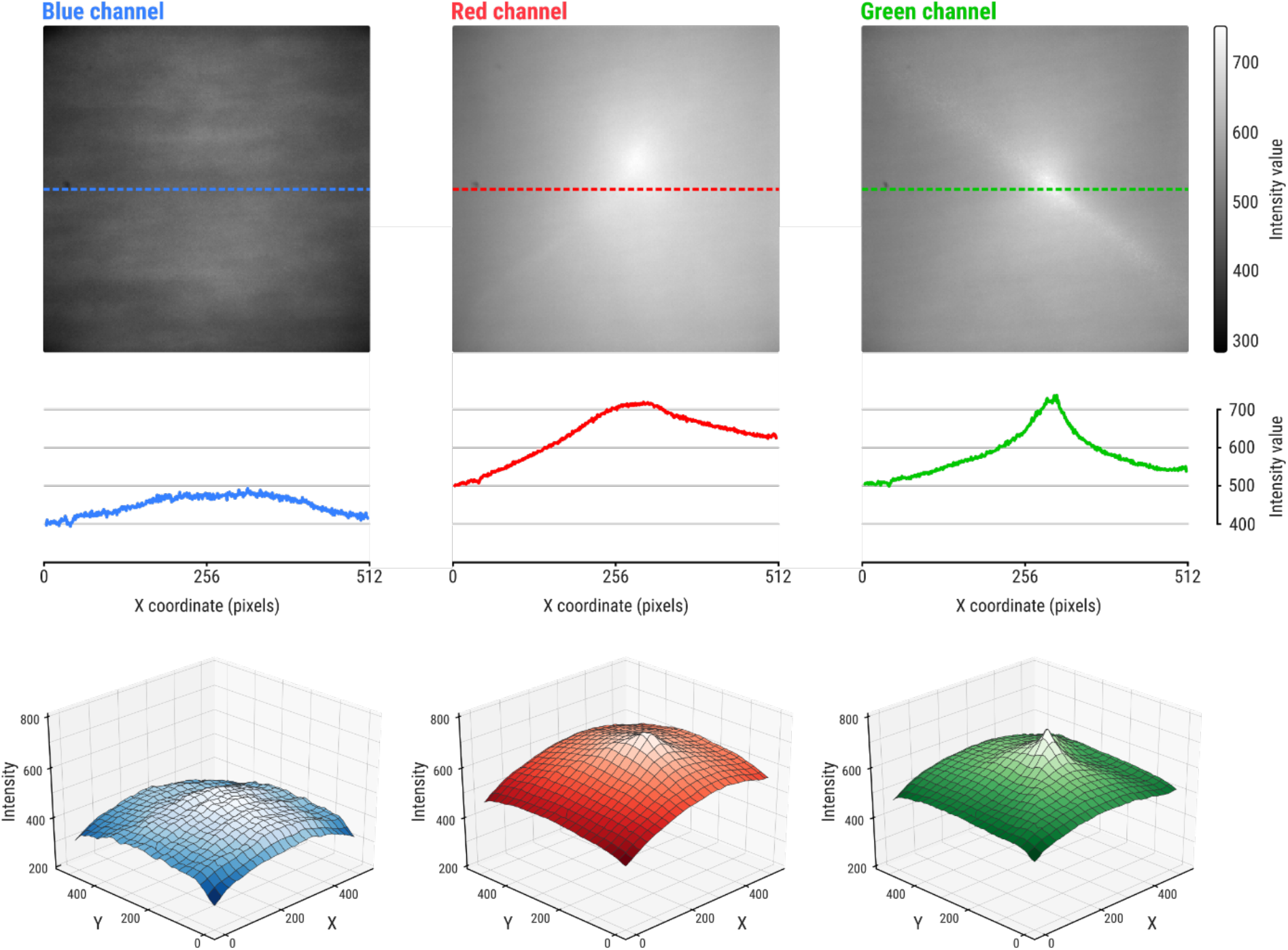
Scan illumination functions. **(Top)** Blue, red and green median illumination functions computed from the 1,140 images of a single scan. The intensity profiles across the middle of the FOV (y=256 px, dashed lines, **Middle**) and the 3D plots of illumination functions **(Bottom)** show roughly similar intensity values ([400; 700] interval on 16-bit images) but different shapes of the three illumination functions.

**Figure S2.**
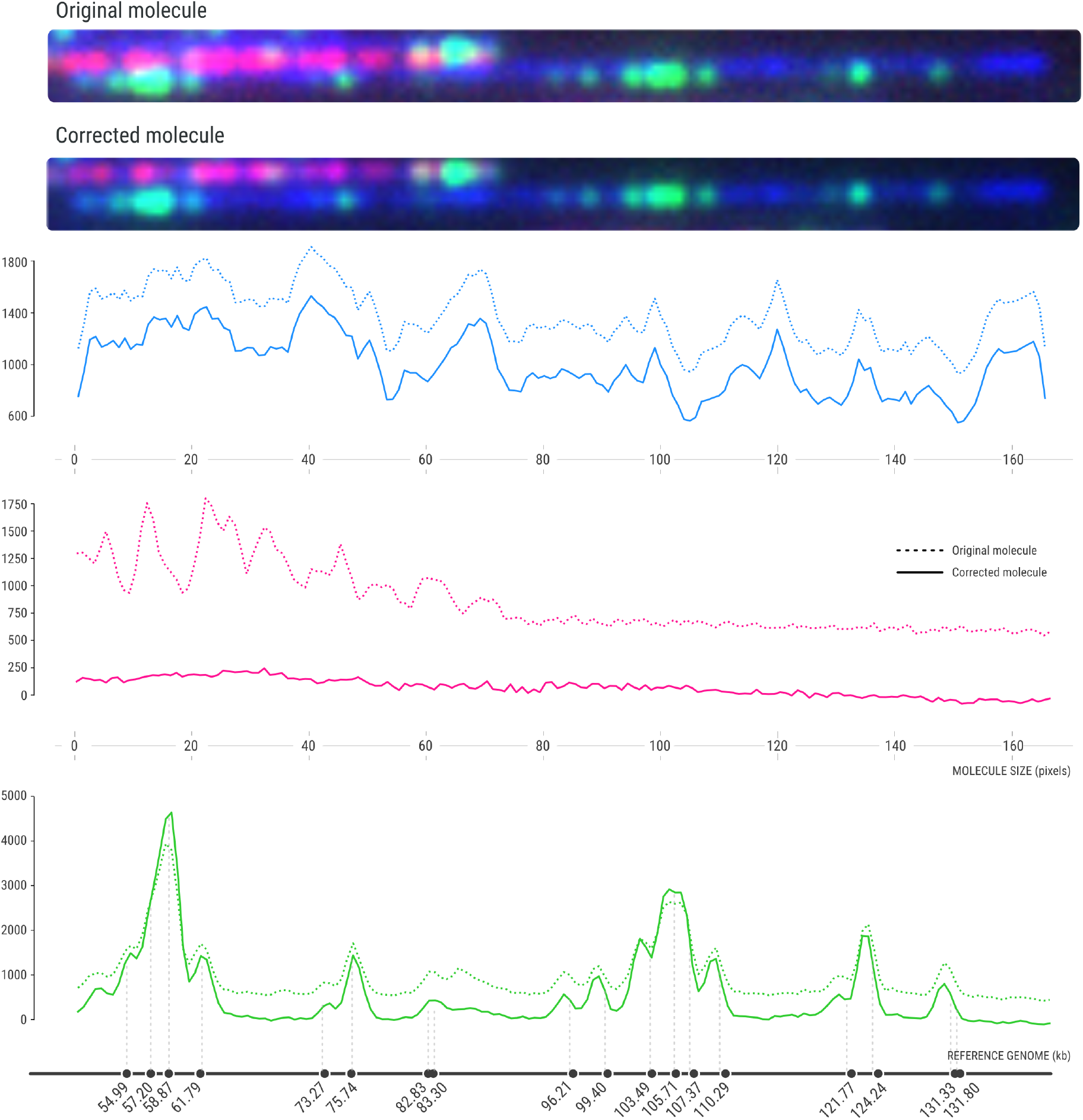
Image correction avoids false signal collection from an adjacent DNA molecule. This Figure is related to Figure 4. The two image crops show two adjacent DNA molecules before and after correction of uneven illumination and chromatic shift. Fluorescent intensity profiles (blue, DNA; red, replication; green, barcodes) of the bottom DNA molecule before (dotted lines) and after (solid lines) image correction are shown. The red signal emitted by the top molecule is erroneously collected onto the bottom molecule profile in the absence of image correction. This problem is eliminated after image correction.

